# Identification of Gli1 as a progenitor cell marker for meniscus development and injury repair

**DOI:** 10.1101/2020.11.27.401463

**Authors:** Yulong Wei, Hao Sun, Tao Gui, Lutian Yao, Leilei Zhong, Wei Yu, Su-Jin Heo, Lin Han, X. Sherry Liu, Yejia Zhang, Eiki Koyama, Fanxin Long, Miltiadis Zgonis, Robert L Mauck, Jaimo Ahn, Ling Qin

**Author notes:** Corresponding author: Ling Qin, Department of Orthopaedic Surgery, Perelman School of Medicine, University of Pennsylvania, 311A Stemmler Hall, 36th Street and Hamilton Walk, Philadelphia, PA 19104, USA. These authors contributed equally to this article (Yulong Wei, Hao Sun, Tao Gui).

## Abstract

Meniscal tears are associated with a high risk of osteoarthritis but currently have no disease-modifying therapies. Using Gli1-CreER tdTomato mice, we found that Gli1+ cells contribute to the development of meniscus horns from 2 weeks of age. In adult mice, Gli1+ cells resided at the superficial layer of meniscus and expressed known mesenchymal progenitor markers. In culture, meniscal Gli1+ cells possessed high progenitor activities under the control of Hh signal. Meniscus injury at the anterior horn induced a quick expansion of Gli1+ cells. Normally, the tissue healed slowly, leading to cartilage degeneration. Ablation of Gli1+ cells further hindered this repair process. Strikingly, intra-articular injection of Gli1+ meniscal cells or an Hh activator right after injury accelerated the bridging of the interrupted ends and attenuated signs of osteoarthritis. Taken together, our work identified a novel progenitor population in meniscus and proposes a new treatment for repairing injured meniscus and preventing osteoarthritis.

## Introduction

Meniscal tears, with all the morbidity and disability they cause, are among the most common injuries of the knee affecting both the young and aged; and the procedures to address them are among the most commonly performed surgeries in the orthopedics field. Beyond the short-term pain, disability, time from desired activities including work, meniscal injuries are important early events in the initiation and later propagation of osteoarthritis (OA) [1]. From a clinical therapeutic point of view, surgical treatments, including the maximally preserving partial meniscectomy, while improving immediate symptoms, do not delay the natural history progression of OA or may actually accelerate it. As the adult meniscus is predominantly avascular, true biologic healing with surgical repair remains a viable treatment for only a small portion of individuals typically with tears contained within the red vascular zone [2]. For the majorities of injuries, a restorative biologic therapy does not currently exist in practice.

Mesenchymal progenitors play a critical role in tissue regeneration. Therefore, identifying and characterizing residential mesenchymal progenitors in meniscus are important for developing novel and effective strategies to treat meniscus injury. Using enzymatic digestion and clonal expansion methods, previous studies have demonstrated that human and rabbit meniscus contain mesenchymal progenitors with multi-differentiation abilities [3-6]. Interestingly, the superficial layer of meniscus was proposed to harbor the progenitors. By collecting cells growing out of mouse meniscus explant, Gamer et al. showed that these cells exhibit stem cell-like characteristic and are located in the superficial zone in vivo [7]. During injury, it has been observed that progenitors on the meniscus surface migrate from vascularized red zone to non-vascularized white zone for repair [8]. While these cells in culture express several common mesenchymal progenitor markers, such as CD44, Sca1, and CD90, their in vivo properties and regulatory signaling pathways are not known [6, 8].

Hedgehog (Hh) signaling is essential for embryonic development and tissue homeostasis. It is one of few fundamental pathways that maintain adult stem and progenitor cells in various organs, such as brain, skin, bladder, teeth, and others [9]. Following injury, Hh signaling can trigger stem and other resident cells to participate in repair, and therefore, Hh upregulation is viewed not only as a natural response to injury but also as a way to stimulate tissue repair by activating stem cells. Gli1, an integral effector protein of Hh pathway, was recently recognized as a marker for bone marrow, periosteal, and periarticular mesenchymal progenitors [10-12], suggesting that Hh signaling is also functional in skeleton for maintaining tissue-specific stem and progenitors.

In this study, we constructed a Hh reporter mouse (*Gli1-CreER Tomato, Gli1ER/Td*), and found that Gli1-labeled Td^+^ cells are exclusively located in the horns of adolescent meniscus. These cells contribute to meniscus development and possess mesenchymal progenitor properties. In adult mice, Gli1^+^ cells mostly reside at the superficial layer of meniscus and they rarely become cells in the center of meniscus. Interestingly, meniscus injury induced a rapid expansion of Gli1^+^ cells and elimination of these cells mitigated repair. Using sorted Gli1-labeled cells and a Gli1 activator, we demonstrated that activating Hh signaling could be an effective way to promote meniscus repair and prevent OA progression.

## Materials and Methods

### Animals

All animal work performed in this report was approved by the Institutional Animal Care and Use Committee (IACUC) at the University of Pennsylvania. *Gli1-CreER Rosa-tdTomato* (*Gli1ER/Td*) mice were generated by breeding *Gli1-CreER* mice (Jackson Laboratory, Bar Harbor, ME USA) with *Rosa-tdTomato* mice (Jackson Laboratory). They were further bred with *Rosa-DTA* mice (Jackson Laboratory) to produce *Gli1-CreER Rosa-tdTomato Rosa-DTA* (*Gli1ER/Td/DTA*). In accordance with the standards for animal housing, mice were group housed at 23-25°C with a 12 h light/dark cycle and allowed free access to water and standard laboratory pellets. All animal work performed in this report was approved by the Institutional Animal Care and Use Committee (IACUC) at the University of Pennsylvania.

To induce Td expression or ablate Gli1-labeled cells, mice (*Gli1ER/Td* or *Gli1ER/Td/DTA*) received vehicle or Tamoxifen (Tam) injections at 50 mg/kg at P4 and P5 or 75 mg/kg for 5 days at ages older than 1 week. For EdU labeling of proliferation experiment, mice were injected with daily 1.6 mg/kg EdU (Invitrogen, Carlsbad, USA, A10044) for 4 days before harvesting. For EdU labeling of slow-cycling experiment, mice were injected with daily 5 mg/kg EdU for 4 days at P3-6.

Male mice at 3 months of age were subjected to meniscus injury at right knees. To perform the surgery, the joint capsule was opened immediately after anesthesia and the anteriomedial horn of meniscus were cut into two parts using microsurgical scissors. The joint capsule and the subcutaneous layer were then closed with suture followed by skin closure with tissue adhesive. In sham surgery, meniscus will be visualized but not transected. For cell treatment, cells digested from meniscus of *Gli1ER/Td* mice were sorted by FACS to collect Td^+^ and Td^-^ cells. 10,000 cells were injected into the knee joint space of sibling *WT* mice immediately after meniscus surgery. For activator treatment, 2 μl purmorphamine (100 μM) were injected into the knee joint space of *WT* or *Gli1ER/Td* mice immediately after surgery. Mice were euthanized at indicated time points for histology analysis.

The knee joint pain after meniscus injury was evaluated in mice at 1 month after surgery using von Frey filaments as described previously [13]. An individual mouse was placed on a wire-mesh platform (Excellent Technology Co.) under a 4×3×7 cm cage to restrict their move. Mice were trained to be accustomed to this condition every day starting from 7 days before the test. During the test, a set of von Frey fibers (Stoelting Touch Test Sensory Evaluator Kit #2 to #9; ranging from 0.015 to 1.3 g force) were applied to the plantar surface of the hind paw until the fibers bowed, and then held for 3 seconds. The threshold force required to elicit withdrawal of the paw (median 50% withdrawal) was determined five times on each hind paw with sequential measurements separated by at least 5 min.

To induce OA, male mice at 3 months of age were subjected to DMM surgery at right knees and sham surgery at left knees as described previously [14]. Briefly, in DMM surgery, the joint capsule was opened immediately after anesthesia and the medial meniscotibial ligament was cut to destabilize the meniscus without damaging other tissues. In sham surgery, the joint capsule was opened in the same fashion but without any further damage.

### Human and Mini-pig Meniscus Samples

The meniscus samples were prepared from the de-identified specimens obtained at the total arthroplasty of the knee joints and used for histological and immunohistochemical examination. The meniscus degeneration severity was evaluated according to the meniscus surface including lamellar layer, cellularity, collagen organization and safranin O/fast green staining [15]. 6-month-old male Yucatan minipigs were utilized (Sinclair Bioresources) to provide meniscus tissues. Anterior horn meniscus tissue was obtained for following histological analysis.

### Histology

After euthanasia, mouse knee joints were harvested and fixed in 4% PFA overnight followed by decalcification in 10% M EDTA (pH 7.4). Samples were processed for either cryosections after 1 week of decalcification or paraffin sections after 4 weeks of decalcification. For healthy knee joints, a serial of 6 μm-thick sections were cut across the entire compartment of the joint at the coronal or sagittal plane followed by fluorescent imaging (cryosections) or safranin O/fast green staining for brightfield imaging (paraffin sections). For meniscus injured knee joints, a serial of 6 μm-thick sections were cut across the entire anterior horn area in the direction perpendicular to the meniscus injury gap (oblique sections) followed by fluorescent imaging (cryosections) or safranin O/fast green staining for brightfield imaging (paraffin sections). To evaluate meniscus healing process, we collected all sections (∼15) including both synovial and ligamental ends.

Three sections were selected from each knee, corresponding to 1/3 (sections 1-5), 2/3 (sections 6-10), and 3/3 (sections 11-15) regions of the entire section set to quantify the meniscus repair scores according to the connection between two ends, existence of fibrochondrocyte and sensitivity of safranin O staining [16]. The method to measure Mankin Score was described previously [17]. Briefly, two sections within every consecutive six sections in the entire sagittal section set for each knee were stained with safranin O/fast green and scored by two blinded observers (YW and HS). Each knee received a single score representing the maximal score of its sections.

For immunohistochemistry staining, mouse, porcine, and human paraffin sections were incubated with rabbit anti-Gli1 (NOVUS biologicals, NB600-600) and anti-Ki67 (Abcam, ab15580) at 4°C overnight followed by binding with biotinylated secondary antibody incubation for 1h and DAB color development. For immunofluorescence staining, sagittal knee joint cryosections from 12-week-old Gli1ER/Td mice were incubated with rat anti-sca1 (Santa cruz, sc-52601), rat anti-Cd200 (Santa cruz, sc-53100), mouse anti-Cd90 (Santa cruz, sc-53456), mouse anti-PDGFRα (Santa cruz, sc-398206), mouse anti-Cd248 (Santa cruz, sc-377221), rabbit anti-Prg4 (Abcam, ab28484) at 4°C overnight followed by binding with corresponding Alexa Fluor® 488-conjugated secondary antibody incubation for 2h and DAPI counterstaining.

### Primary Mouse Meniscus Cell Culture

Mouse menisci were dissected from tibiae of 4-week-old mice and digested in 0.25% Trypsin-EDTA (Gibco) for 1 h followed by 300U/mL collagenase type I (Worthington Biochemical) for 2 h. Cells from the second digestion were cultured in the growth medium (αMEM supplemented with 10% fetal bovine serum plus 100 IU/ mL penicillin and 100 mg/mL streptomycin) to obtain meniscus cell culture. For CFU-F assay, digested cells were seeded at 20,000 cells per T25 flask. Seven days later, flasks were stained with 3% crystal violet to quantify colony numbers. To study cell migration, primary meniscus cells were seeded in 12-well plates. When reaching confluency, the cell layer was scratched by a 1000 μL pipette tip and then cultured in FBS free growth medium. Wound closure was monitored by imaging at 0 and 48 hr later. To study cell proliferation, primary meniscus cells were seeded at 50,000 cells/well in 12-well plates and cell numbers were counted 2, 4, and 6 days later.

Chondrogenic, osteogenic, and adipogenic differentiation was performed as describe previously [12]. For meniscal differentiation, confluent cells were cultured in meniscus differentiation medium (high glucose DMEM with 100 IU/ mL penicillin, 100 mg/mL streptomycin, 0.1 μM dexamethasone, 50 μg/ml ascorbate 2-phosphate, 40 μg/ml l-proline, 100 μg/ml sodium pyruvate, 6.25 μg/ml insulin, 6.25 μg/ml transferrin, 6.25 ng/ml selenous acid, 1.25 mg/ml bovine serum albumin, 5.35 μg/ml linoleic acid and 10 ng/ml TGF-β3) as described previously [18, 19].

### Flow Cytometry and Cell Sorting

Flow cytometry and cell sorting were performed on a FACS Aria III cell sorter (BD Biosciences) and analyzed using Flow Jo software (Tree Star). Digested meniscus cells were re-suspended in flow buffer (2% FBS/PBS) and stained with Sca1 (BioLegend, 108131), Cd90 (BioLegend, 202526), Cd200 (BioLegend, 123809), and PDGFRα (BioLegend, 135907) flow antibody for 1 h at 4°C. After PBS wash, cells were analyzed by flow cytometry or sorted for Gli1^+^ and Gli1^-^ cells.

### RNA Analyses

To quantify the expression level of marker genes, total RNA was collected in Tri Reagent (Sigma, St. Louis, MO, USA) for RNA purification. A Taqman Reverse Transcription Kit (Applied BioSystems, Inc., Foster City, CA, USA) was used to reverse transcribe mRNA into cDNA. The power SYBR Green PCR Master Mix Kit (Applied BioSystems, Inc) was used for quantitative real-time PCR (qRT-PCR). The primer sequences for the genes used in this study are listed in Supplemental Table S1.

### Statistical Analyses

Data are expressed as means ± standard error of the mean (SEM) and analyzed by t-tests, one-way ANOVA with Dunnett’s or Turkey’s posttest and two-way ANOVA with Turkey’s post-test for multiple comparisons using Prism 8 software (GraphPad Software, San Diego, CA). For assays using primary cells, experiments were repeated independently at least three times and representative data were shown here. Values of p<0.05 were considered statistically significant.

## Results

### The expression patterns of Gli1 ^+^ cells and their descendants in mouse meniscus

We performed lineage tracing with *Gli1ER/Td* mice at various ages to identify Gli1^+^ cells and their descendants at 6 weeks later in meniscus (Fig. S1A). Joints were cut at either sagittal or coronal planes to visualize different parts of meniscus (Fig. S1B). In line with our previous report [11], at 1 week of age, Gli1^+^ cells were only observed in the periarticular layer of articular cartilage, but not in the meniscus and other joint tissues (Fig. 1Aa-c). Long term tracing also did not detect any Td signal in the meniscus, confirming that neonatal meniscus does not harbor Gli1^+^ cells (Fig. 1Ad). At 2 weeks of age, most cells in the anterior horn of the meniscus, both medially and laterally, were Td^+^ (Fig. 1Ae-g). Six weeks of tracing confirmed that the entire anterior horn, but not the posterior horn, is labeled by Td (Fig. 1Ah). At 4 weeks of age, Gli1^+^ cells were concentrated in the superficial layer of the anterior horn; 6 weeks later, most cells in both superficial and central portions of the anterior horn were labeled by Td (Fig. 1Ai-l). Within the posterior horn, very few cells in the center of meniscus were initially labeled but then gave rise to the majority of internal cells 6 weeks later. Quantification along the length of the meniscus over the time indicated that 1-8 weeks of age represents the rapid growing phase for the meniscus (Fig. S2). Taken together, our data suggested that Gli1^+^ cells represent progenitors for meniscus cells of the horn regions at adolescence stage.

**Figure 1.**
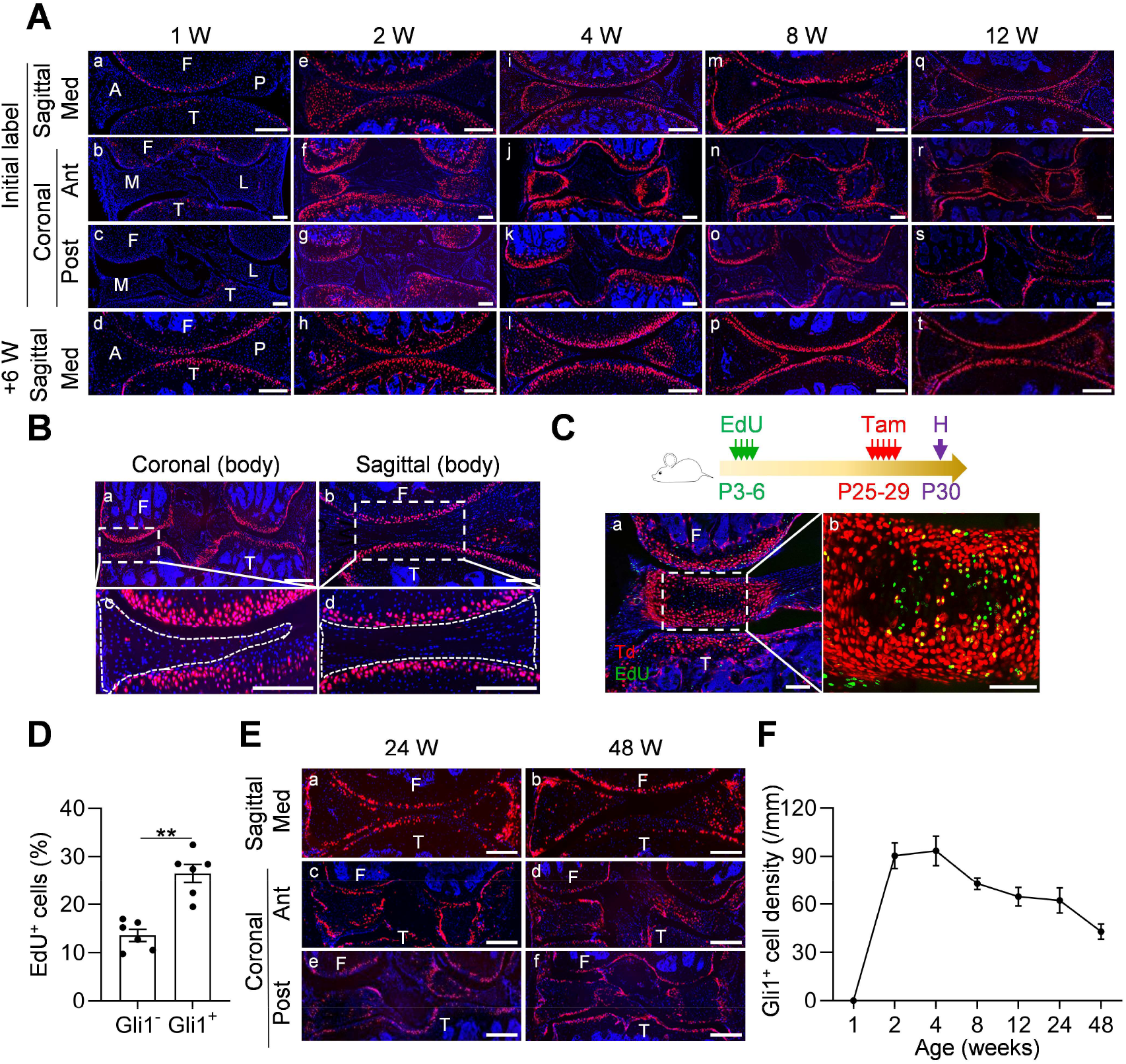
Gli1 labels mesenchymal progenitors in mouse meniscus during development. (**A**) Representative fluorescence images of meniscus sections at indicated ages and sectioning sites. n = 3 mice/age/sectioning site. Scale bars, 200 μm. F: femur; T: tibia; A: anterior; P: posterior; M: medial meniscus; L: lateral meniscus; Med: medial; Ant: anterior; Post: posterior; Red: Td; Blue: DAPI. (**B**) Representative fluorescence images of meniscus body at coronal (a) and sagittal (b) planes from 12-week-old *Gli1ER/Td* mice. Meniscus were harvested at 24 h after last Tam injection. n = 3 mice/sectioning site. Scale bars, 200 μm. Boxed areas in a and b are shown at high magnification as c and d, respectively. Dashed lines outline meniscus. F: femur, T: tibia; Red: Td; Blue: DAPI. (**C**) Top panel is a schematic representation of the study protocol. *Gli1ER/Td* mice were injected with EdU at P3-6 and Tam at P25-29. Joints were harvested 24 h later. Representative confocal images of coronal sections of mouse knee joints are presented at the bottom panel. Boxed area in a (Scale bars, 200 μm) is shown at high magnification in b (Scale bars, 50 μm). F: femur, T: tibia; Red: Td; Blue: DAPI; Green: EdU. (**D**) The percentage of EdU^+^ cells within Gli1^+^ or Gli1^-^ meniscus cells were quantified. n = 6 mice/group. (**E**) *Gli1ER/Td* mice were treated with Tam at 24 or 48 weeks of age and analyzed 24 h later. Representative fluorescence images of sagittal (a, b) and coronal (c-f) sections of knee joints are presented. Scale bars, 200 μm. F: femur, T: tibia; Red: Td; Blue: DAPI. (**F**) The density of Gli1^+^ cells along meniscus surface was measured in mice at different ages. n = 5 mice/age. Statistical analysis was performed using unpaired two-tailed t-test. Data presented as mean ± s.e.m. **p < 0.01.

Starting from 8 weeks of age, Gli1^+^ cells were exclusively restricted to the superficial layer of the anterior horn throughout tracing (Fig. 1Am-t). The labeling pattern in the posterior horn was slightly different that Td^+^ cells first appear in the center and then expand to the entire tissue at 6 weeks later (Fig. 1Am-p). At 12 weeks of age, Gli1^+^ cells remained restricted to the superficial layer of both anterior and posterior horns throughout tracing (Fig.1Aq-t). At any given age, Td signal was not detected in the center of the body of either the medial or lateral meniscus regardless of cutting planes (Fig. 1B).

Slow cycling cells are considered quiescent stem cells [20]. Applying a label-retention method on neonatal mice (EdU injections at P3-6 and Tam injections at P25-29), we found that Gli1-labeled cells at P30 contain much more EdU^+^ cells than non-Gli1-labeled cells (Fig. 1C, D), indicating that meniscus stem cells are enriched in the Gli1^+^ cell population.

When mice reached mature and late adult stages (24 and 48 weeks of age, respectively), Gli1 mostly marked the superficial layer of both anterior and posterior horn of the meniscus (Fig. 1E). Quantification of cells along the surface of meniscal horns revealed a drastic reduction of Gli1^+^ cells in aged mice compared to adolescent mice (Fig. 1F).

Meniscus is attached to neighboring bones via fibrocartilaginous entheses. We found that Gli1 labels these entheses between anteromedial, posteromedial, anterolateral, posterolateral meniscus and the tibial plateau or femur condyle (Fig. S3A). In addition, Gli1 also labeled the osseous ligamentous junctions between the anterior cruciate ligament or posterior cruciate ligament and femur or tibia (Fig. S3B, C).

The existence of Gli1-labeled cells on the meniscus surface was confirmed by Gli1 immunostaining (Fig. S4A). Furthermore, analysis of porcine meniscus revealed a similar staining pattern. As shown in Fig. S4B, Gli1^+^ cells were located in the superficial layer, but not in the central part, of meniscus horn in the adult mini-pig.

### Gli1-expressing meniscus cells are mesenchymal progenitors

We next investigated whether Gli1^+^ meniscus cells possess mesenchymal progenitor properties. Immunostaining of mesenchymal markers, such as Sca1, Cd90, Cd200, PDGFRα [21] and Cd248 [22, 23] revealed their co-staining with Td^+^ signal in the superficial layer of the meniscus in 3-month-old mice (Fig. 2Aa-e). Prg4 is the lubricant highly synthesized by cartilage, meniscus and synovium surface cells [13, 24]. We found that Gli1^+^ cells are also Prg4^+^ (Fig. 2Af). Using an enzymatic digestion approach, we harvested meniscus cells for subsequent studies. Flow cytometry revealed that Gli1^+^ cells are only 2% of meniscus cells digested from 3-month-old mice and that they express mesenchymal progenitor markers Sca1, CD90, CD200, and PDGFRα at a higher level than Gli1^-^ cells (Fig. S5, Fig. 2B).

**Figure 2.**
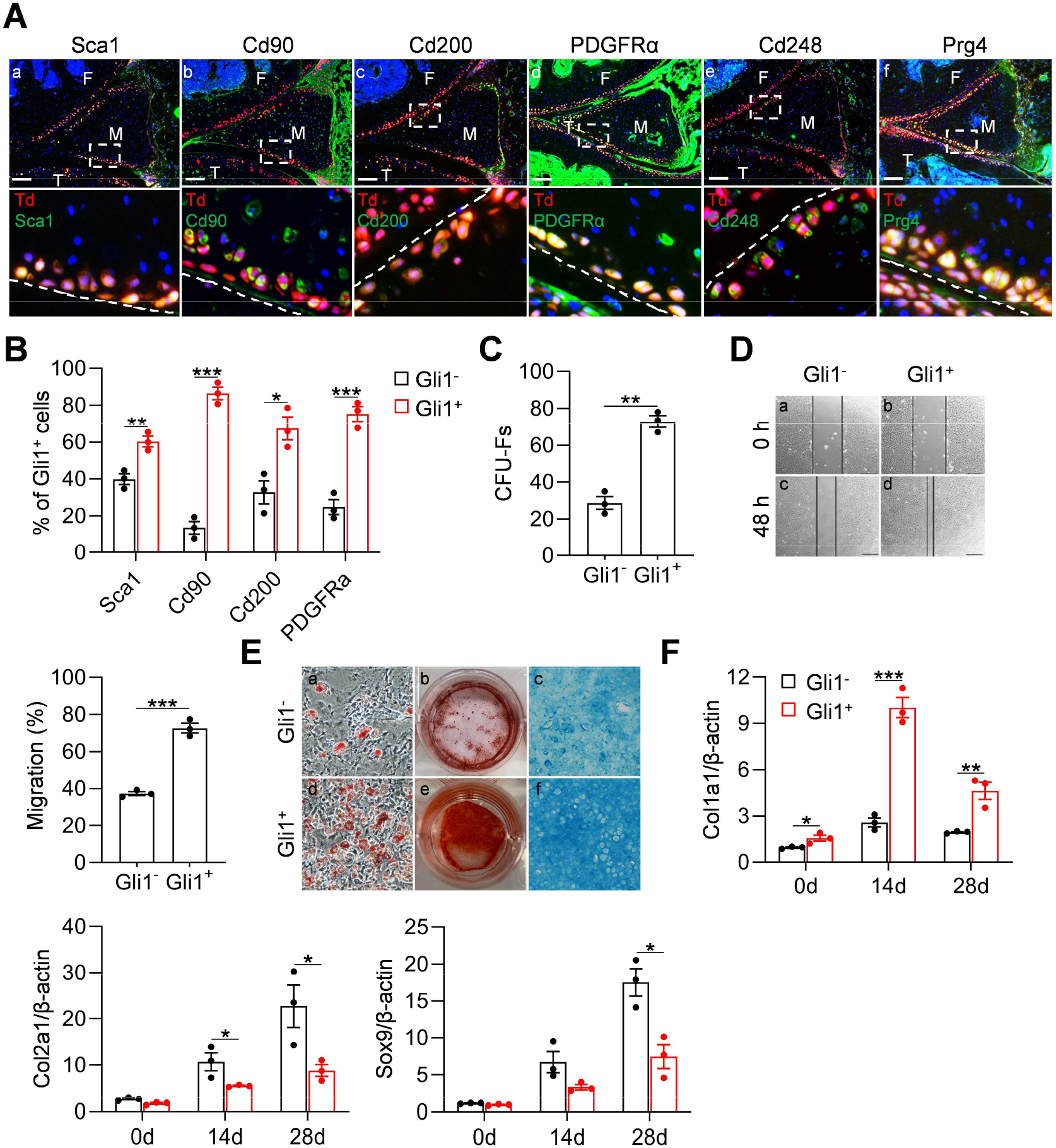
Gli1-labeled meniscus cells possess mesenchymal progenitor properties. (**A**) Representative immunofluorescence images of Sca1, Cd90, Cd200, PDGFRα, Cd248 and Prg4 in 3-month-old *Gli1ER/Td* meniscus. Scale bars, 200 μm. Boxed areas are shown at high magnification in corresponding panels to the bottom. Dashed lines indicate the surface of meniscus. Yellow cells are double positive for progenitor marker and Td. Blue: DAPI; F: femur; T: tibia; M: meniscus. (**B**) Quantification of the expression level of mesenchymal progenitor markers in Gli1^+^ and Gli1^-^ cells from meniscus. Digested meniscus cells from 3-month-old *Gli1ER/Td* mice were subjected to flow cytometry analysis. n = 3 independent experiments. (**C**) CFU-F assay of Gli1^+^ and Gli1^-^ cells sorted by FACS from digested meniscus cells. n = 3 independent experiments. (**D**) Representative bright-field images of the scratch-wound closure in Gli1^+^ or Gli1^-^ meniscus cells. Scale bars, 200 μm. Solid lines indicate the remaining area not covered by meniscus cells. The relative migration rate was measured by the percentage of scratched area being covered by migrated cells at 48 h. n = 3 independent experiments. (**E**) Representative adipogenic (AD), osteogenic (OB), and chondrogenic (CH) differentiation images of Gli1^+^ and Gli1^-^ cells. Cells were stained by Oil Red, Alizarin red, and Alcian blue, respectively. (**F**) qRT-PCR analysis of Col1a1, Col2a1 and Sox9 mRNA in Gli1^+^ and Gli1^-^ cells at day 0, 14, and 28 of meniscal differentiation. n = 3 independent experiments. Statistical analysis was performed using unpaired two-tailed t-test. Data presented as mean ± s.e.m. *p < 0.05, **p < 0.01, ***p < 0.001.

Digested meniscus cells formed CFU-Fs in dishes. Interestingly, 72.8% of CFU-Fs were Td^+^, suggesting that Gli1^+^ cells have a high clonogenic activity (Fig. 2C). While both Td^+^ and Td^-^ cell can grow in culture, sorted Td^+^ cells proliferated faster (Fig. S6A, B) and migrated quicker (Fig. 2D) than Td^-^ cells. In addition, Td^+^ meniscus cells had a better ability to undergo osteogenic, adipogenic, and chondrogenic differentiation than Td^-^ cells in vitro (Fig. 2E). When subjected to meniscal differentiation, Td^+^ cells expressed much more *Col1a1*, a meniscal marker, and less *Col2a1* and *Sox9*, two chondrogenic markers, than Td^-^ cells (Fig. 2F). Taken together, these data demonstrated that Gli1^+^ cells possess the properties of mesenchymal progenitors: self-renewal and multi-lineage differentiation.

### Hh signaling regulates cell behavior of meniscus progenitors

Gli1 expression is a reporter for Hh signaling pathway [25]. To investigate whether Hh signaling is involved in regulating meniscus progenitors, we treated meniscus progenitors with purmorphamine, an activator of Hh signaling [26], or Gli antagonist 61 (GANT-61), a Gli1 inhibitor [27], and performed proliferation, migration, and differentiation assays. In line with previous reports [28, 29], GANT-61 reduced Gli1 expression in meniscus progenitors while purmorphamine increased it (Fig. 3A). Cell counting revealed that GANT-61 reduces the number of meniscus cells in culture over time while purmorphamine increases cell number (Fig. 3B). Similarly, scratch assay showed that Hh signaling activation is required for the migration of meniscus progenitors (Fig. 3C, D). Furthermore, as shown by qRT-PCR, inhibition of Gli1 by GANT-61 reduces *Col1a1, Col2a1*, and *Sox9* expression and activation of Gli1 by purmorphamine has the opposite effects during meniscus differentiation (Fig. 3E). To further support the above proliferation data, purmorphamine up-regulated the expression of cell cycle gene *Ccnd1* and down-regulates the expression of cell cycle inhibitor *Cdkn2a*, while GANT-61 stimulates the expression of *Cdkn2a* (Fig. 3E). Our data demonstrated an important action of Hh signaling in promoting proliferation, migration, and differentiation of meniscus progenitors.

**Figure 3.**
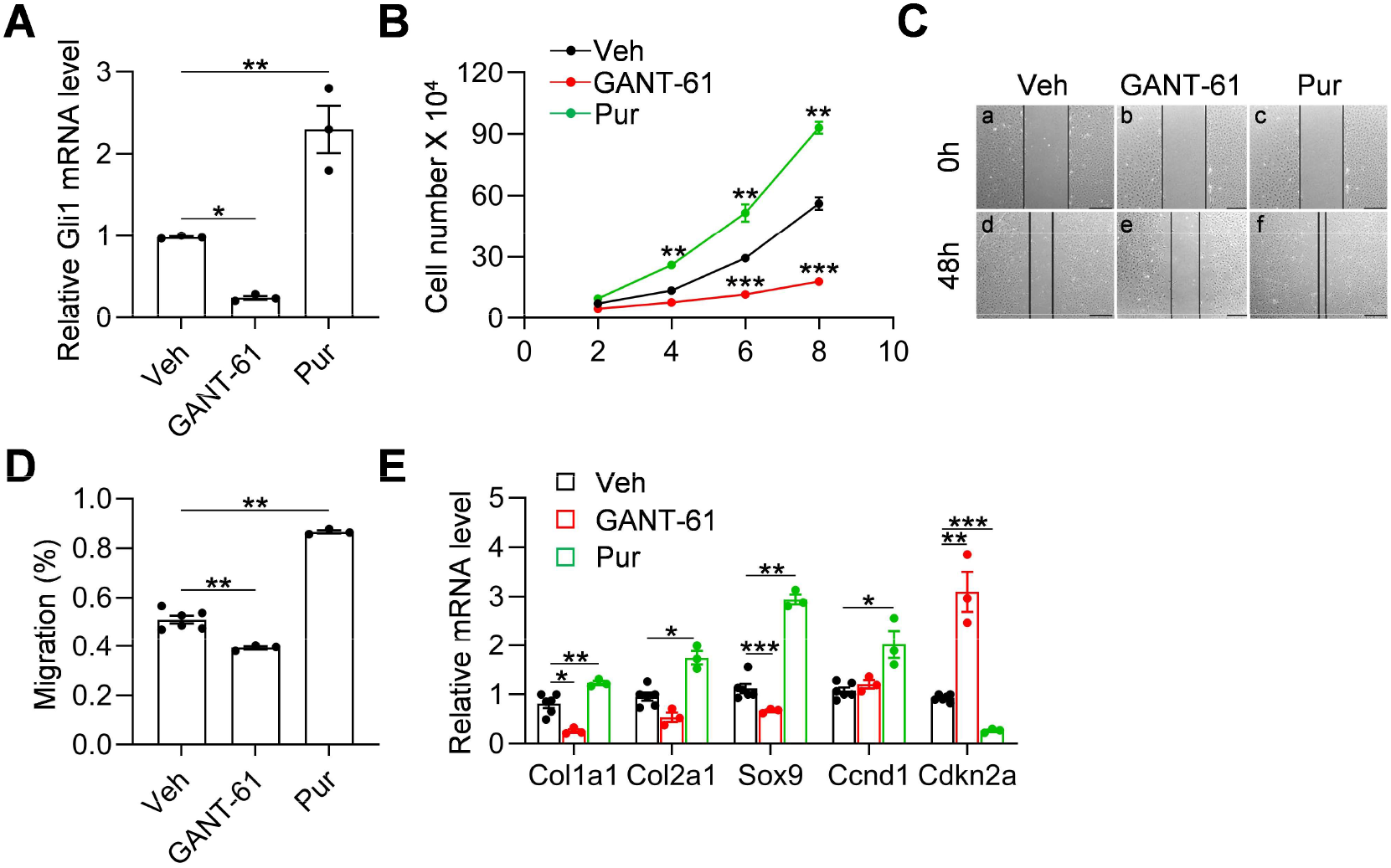
Hh signaling stimulates proliferation and migration of meniscus progenitors. (**A**) qRT-PCR analysis of Gli1 mRNA in primary mouse meniscus cells treated with vehicle, GANT-61 or purmorphamine (Pur) for 48 h. n = 3 independent experiments. (**B**) The proliferative ability of primary mouse meniscus cells was up-regulated by purmorphamine and down-regulated by GANT-61 over 8 days of culture. n = 3 independent experiments. (**C**) Representative bright-field images of the scratch-wound closure in meniscus cells treated with veh, GANT-61 or purmorphamine. Scale bars, 200 μm. Solid lines indicate the remaining area not covered by meniscus cells. (**D**) The relative migration rate was measured. n = 3-6 independent experiments. (**E**) qRT-PCR analysis of marker genes in meniscus cells undergoing meniscal differentiation in the presence or absence of GANT-61 and purmorphamine. n = 3-6 independent experiments. Statistical analysis was performed using one-way ANOVA with Dunnett’s post hoc test. Data presented as mean ± s.e.m. *p < 0.05, **p < 0.01, ***p < 0.001.

### Injury-induced Gli1 ^+^ cell expansion are critical for meniscus healing

Meniscal tear is a common injury in joints. To mimic this injury, we surgically cut the anteromedial horn of the meniscus into two parts in 3-month-old mice, resulting in disconnected synovial and ligamental ends of the meniscus (Fig. S7A). *Gli1ER/Td* mice received Tam right before surgery (Fig. S7B). At 1-2 weeks post-surgery, the two ends of the meniscus retracted toward the synovium and ligament, respectively (Fig. 4Aa-c). This was accompanied by massive synovial hyperplasia that wrapped around the ends of the meniscus and likely stabilized them. At 4 weeks, the synovium returned to relatively normal thickness and the two cut ends of the meniscus were aligned but not connected (Fig. 4Ad). Over time, the connection between the two ends gradually moved toward re-establishment but never reached the normal level even after 3 months post surgery (Fig. 4Ae,f). The meniscus repair scores summarized this trend (Fig. 4B), suggesting that meniscus heals slowly in this injury model.

**Figure 4.**
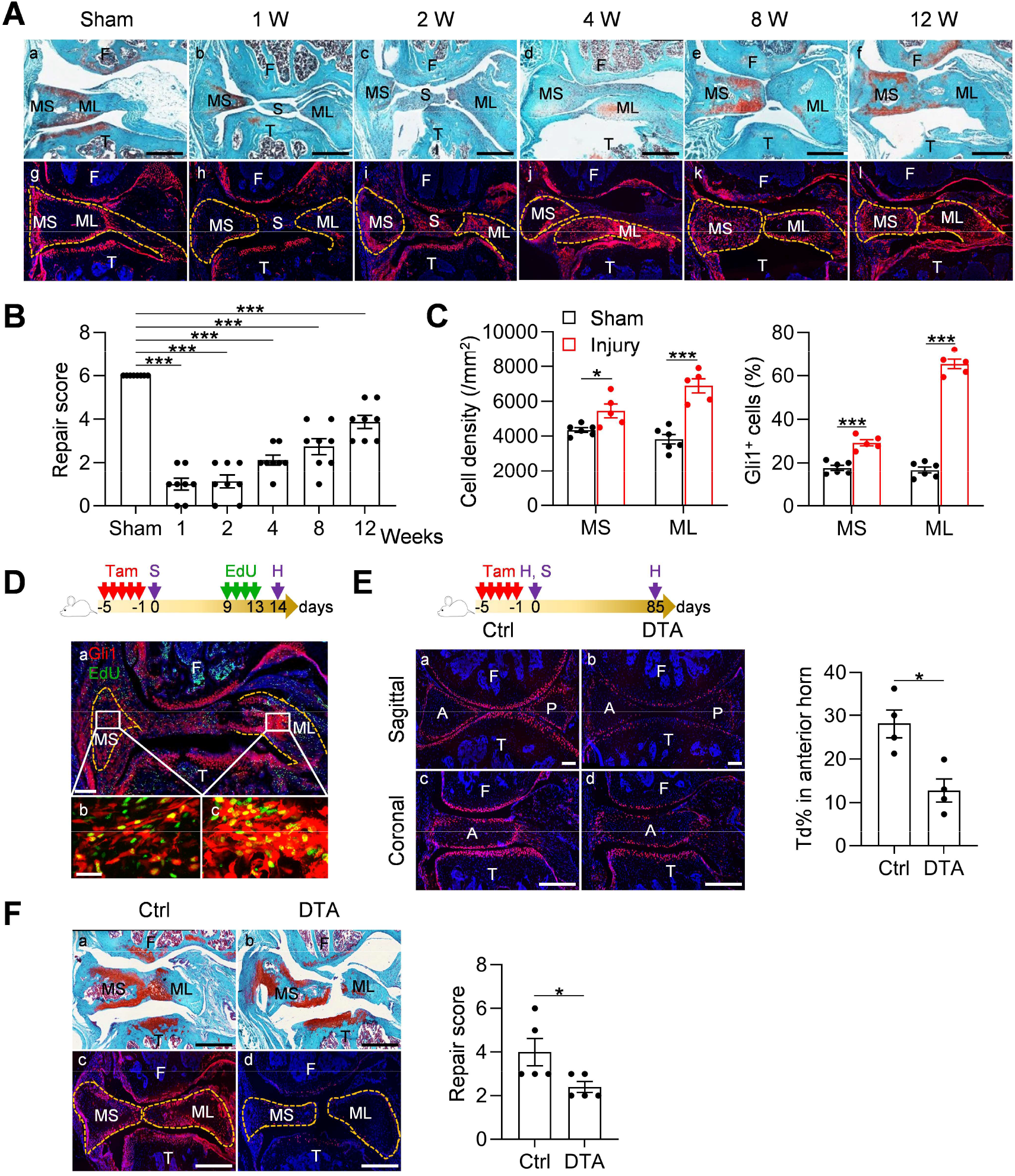
Meniscus injury rapidly expands Gli1-labeled cells. (**A**) Representative safranin O/fast green staining (top) and fluorescence images (bottom) of oblique sections of mouse knee joints harvested at indicated time points after injury. Dashed lines outline the meniscus. Scale bars, 200 μm. F: femur; T: tibia; S: synovium; MS: meniscus synovial end; ML: meniscus ligamental end; Red: Td; Blue: DAPI. (**B**) Repair score was evaluated at indicated time points after meniscus injury. n = 8 mice/group. (**C**) Cell density in the synovial and ligamental ends of meniscus was quantified at 4 weeks post meniscus injury. n = 5-6 mice/group. The percentage of Gli1^+^ cells in the synovial and ligamental ends of meniscus was quantified at 4 weeks post meniscus injury. n = 5-6 mice/group. (**D**) Top panel is a schematic representation of the study protocol. *Gli1ER/Td* mice were treated with Tam and meniscus injury at 12 weeks of age (day 0) followed by EdU injections at day 9-13 and analyzed at 24 h after the last EdU dosing. A representative confocal image of knee joint is shown below (Scale bars, 250 μm). Boxed areas of synovial and ligamental ends of meniscus (MS and ML, respectively) are shown at high magnification at the bottom (Scale bar, 25 μm). F: femur; T: tibia; Red: Td; Blue: DAPI; Green: EdU. (**E**) Top panel is a schematic representation of the study protocol. *Gli1ER/Td* (Ctrl) or *Gli1ER/Td/DTA* (DTA) mice received Tam injections at 12 weeks of age (day 0). Non-injuried knee joints were harvested at 24h after the last Tam dosing (day 1). Injuried knee joints received meniscus surgery at 24h after the last Tam dosing (day 1) and were harvested 3 months later. A representative fluorescent images of sagittal and coronal mouse knee joint sections at 12 weeks of age are shown below (Scale bars, 200 μm). F: femur; T: tibia; A: anterior; P: posterior; Red: Td; Blue: DAPI. Td^+^ cell percentage in the anterior horn was quantified based on the sagittal images. n = 4 mice/group. (**F**) Representative safranin O/fast green staining (top) and fluorescence images (bottom) of oblique sections of mouse knee joints harvested at 12 weeks after injury. Dashed lines outline the meniscus. Scale bars, 200 μm. F: femur; T: tibia; S: synovium; MS: meniscus synovial end; ML: meniscus ligamental end; Red: Td; Blue: DAPI. Repair score was evaluated. n = 5 mice/group. Statistical analysis was performed using one-way ANOVA with Dunnett’s post hoc test for (B) and unpaired two-tailed t-test for (C), (E) and (F). Data presented as mean ± s.e.m. *p < 0.05, ***p < 0.001.

Fluorescence imaging was used to analyze the contribution of Gli1^+^ cells and their descendants during this process. Strikingly, starting from 2 weeks post-injury, Td^+^ cells appeared at the synovial ends and ligamental ends of injured meniscus (Fig. 4Ai). Their number peaked around 4 weeks, and gradually declined thereafter (Fig. 4Aj-l). Total cell density and the percentage of Gli1^+^ cells at both ends were significantly increased after injury, particularly at the ligamental end (Fig. 4C). EdU incorporation experiment confirmed that many Gli1^+^ cells and their progenies are proliferative at 2 weeks post surgery (Fig. 4D). In old mice (52 weeks of age), this expansion of Td^+^ cells after injury was remarkably attenuated and the end-to-end reconnection was much less with a lower repair score than young adult mice 4 weeks later (Fig. S8A, B), indicating that aging diminishes the repair ability of meniscus.

To further understand the role of Gli1^+^ cell expansion in meniscus repair, we generated *Gli1-CreER Tomato DTA* (*Gli1ER/Td/DTA*) mice for a cell ablation experiment. These mice at 3 months of age received Tam injections followed by meniscus injury (Fig. 4E). One day after the Tam injections, Td^+^ cells in meniscus were drastically decreased by 54.5%, as shown by both sagittal and coronal views of meniscus horns (Fig. 4E). Three months later, while two meniscus ends loosely reconnected in vehicle-treated mice, those in Tam-treated mice were still well separated, leading to a significant reduction of repair score (Fig. 4F). Fluorescence imaging confirmed no more expansion of Td^+^ cells in Tam-treated mice. These data clearly indicate an essential role of Gli1^+^ cells in meniscal healing.

To validate our mouse data, we collected healthy and degenerated human meniscus for immunohistochemistry analysis. Degenerated meniscus had surface disruption, collagen fibers disorganization, and positive safranin O/fast green staining as previously reported [15]. Healthy meniscus did not show Gli1 staining (Fig. S9). However, in moderate and severe degenerated meniscus, Gli1 was readily detectable in cell clusters formed in various sizes and characterized by Ki67^+^ staining, indicating that Gli1^+^ cells are proliferative. These results confirmed an expansion of Gli1^+^ cells in human meniscus tissues and a potential action of Hh signaling in human meniscus repair.

### Activation of Hh/Gli1 pathway accelerates mouse meniscus repair

Since Gli1^+^ cells and their descendants were greatly expanded at the early phase of meniscus injury repair, we hypothesized that activation of Hh/Gli1 pathway could stimulate the repair process. We adopted two approaches to test this hypothesis. One was to inject Gli1^+^ cells freshly isolated from *Gli1ER/Td* meniscus into injured knees (Fig.5A). Strikingly, a single injection of cells right after injury resulted in a reconnection of the synovial and ligamental ends of injured meniscus at 4 weeks, leading to a repair score of 4.8 (Fig. 5A, B). At the same time, these two meniscus ends were well separated in both vehicle and Gli1^-^ meniscus cell-treated groups, with a repair score of only 1.8 and 1.9, respectively. Polarizing images clearly showed a disconnection of collagen fibers in mice that received either vehicle or Gli1^-^ cells. However, in Gli1^+^ cell-treated mice, collagen fibers crossed the broken ends of the meniscus, suggesting that the repair does occur at the structural level. Fluorescence imaging revealed that injected Gli1^+^ cells expand and contribute to the newly formed connection at the injury site (Fig. 5C).

**Figure 5.**
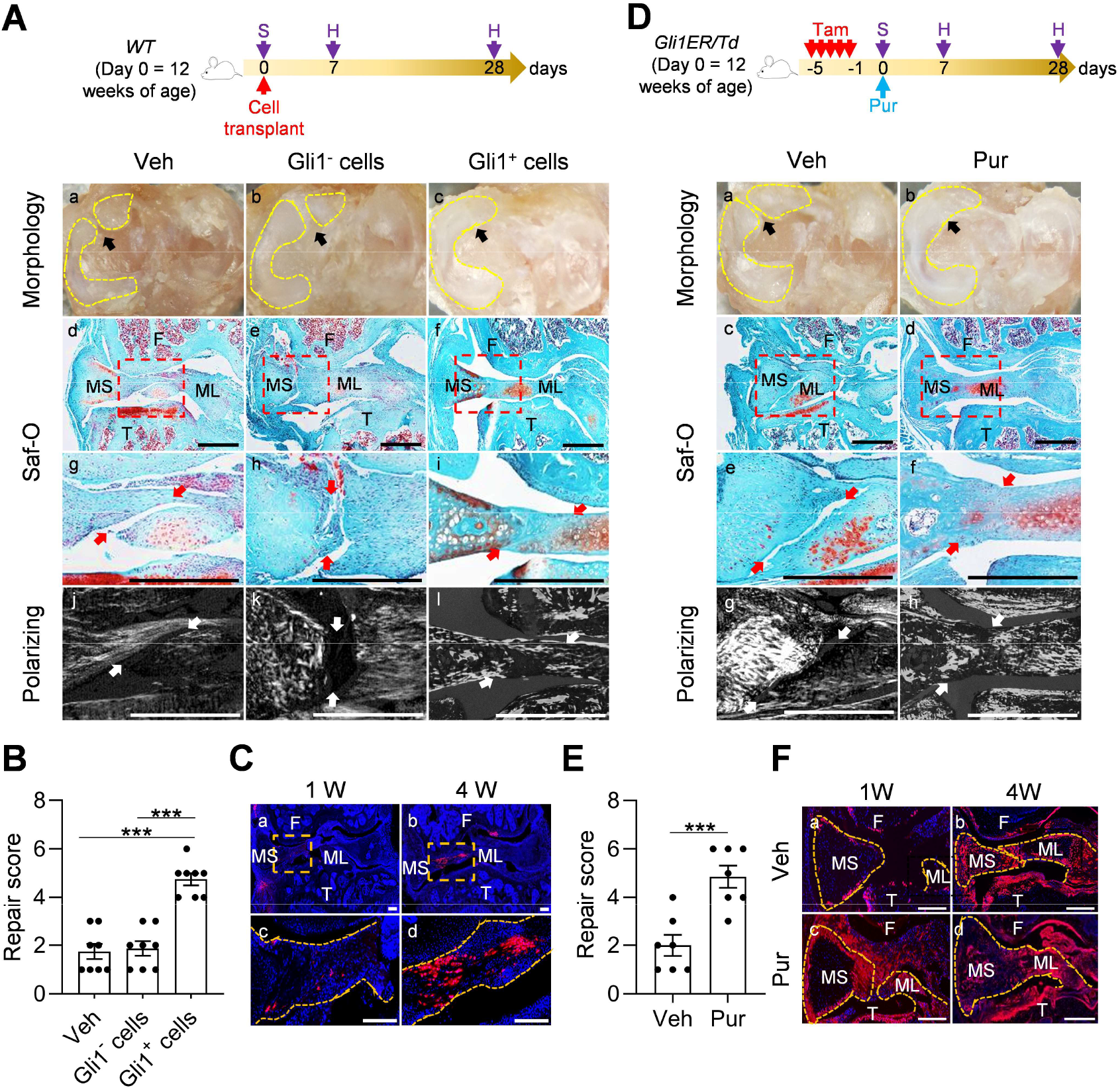
Activation of Hh/Gli1 pathway accelerates mouse meniscus repair. (**A**) Schematic representation of the study protocol. *WT* mice received meniscus injury at 12 weeks of age followed by transplantation of 10,000 Gli1^+^ or Gli1^-^ meniscus cells at the injury site. Knee joints were harvested at 1 and 4 weeks after injury. Representative overview, safranin O/fast green staining, and polarizing images of mouse knee joints at 4 weeks after injury. Yellow dashed lines in a, e, and i outline the overview morphology of injured meniscus. Meniscus is shown attached to tibial plateau. Arrows point to the injury site. Red boxed areas in b, f, and j are shown at high magnification in c, g, and k, respectively. Scale bars, 200 μm. F: femur; T: tibia; MS: meniscus synovial end; ML: meniscus ligamental end. (**B**) Repair score was evaluated. n = 8 mice/group. (**C**) Representative confocal images of mouse knee joints at 1 and 4 weeks after injury and injection of Gli1^+^ cells. Boxed areas in the top panel are shown at a high magnification in the bottom panel. Dashed line outlines meniscus. Scale bars, 200 μm. F: femur; T: tibia; MS: meniscus synovial end; ML: meniscus ligamental end; Blue: DAPI, Red: Td. (**D**) Schematic representation of the study protocol. *Gli1ER/Td* mice received Tam injections and meniscus injury at 12 weeks of age (day 0) followed by vehicle and purmorphamine (pur) injection. Knee joints were harvested at 1 and 4 weeks after injury. Representative overview, safranin O/fast green staining, and polarizing images of mouse knee joints at 4 weeks after injury. Yellow dashed lines in a and e outline the overview morphology of injured meniscus. Meniscus is shown attached to tibial plateau. Red boxed areas in b and f are shown at a high magnification in c and g, respectively. Arrows point to the injury site. Scale bars, 200 μm. F: femur; T: tibia; MS: meniscus synovial end; ML: meniscus ligamental end. (**E**) Repair score was evaluated. n = 7 mice/group. (**F**) Representative fluorescence images of vehicle- and purmorphamine-treated mouse meniscus at 1 and 4 weeks after injury. Scale bars, 200 μm. F: femur; T: tibia; MS: meniscus synovial end; ML: meniscus ligamental end; Blue: DAPI, Red: Td. Statistical analysis was performed using one-way ANOVA with Turkey’s post hoc test for (B) and unpaired two-tailed t-test for (E). Data presented as mean ± s.e.m. ***p < 0.001.

In another approach, we injected purmorphamine to the knee joint right after injury (Fig. 5D). Four weeks later, the injured ends of purmorphamine-treated meniscus were reconnected based on gross morphology, safranin O/fast green staining, and imaging of collagen fibers (Fig. 5D), leading to a repair score of 4.9 (Fig. 5E). There were more Td^+^ meniscus cells in purmorphamine-treated joints than vehicle-treated joints at 1 week after injury (Fig. 5F). These data clearly indicated a therapeutic effect of activating Hh signaling.

### Meniscus repair by enhancing Hh/Gli1 pathway delays OA progression

Meniscal injury inevitably leads to OA in human. To mimic this in mice, we characterized articular cartilage phenotype at 8 weeks post injury. Similar to the surgical destabilization of the medial meniscus (DMM) model of OA, our meniscus injury model caused cartilage degeneration, such as partially loss of proteoglycan, surface fibrillation, and reduction in uncalcified cartilage thickness (Fig. 6A, B). Meanwhile, the calcified cartilage layer was not eroded (Fig. 6B), suggesting a moderate OA with a Mankin Score of 6.9 (Fig. 6C). Strikingly, injections of either Gli1^+^ cells or purmorphamine greatly reduced cartilage degeneration by retaining proteoglycan content, cartilage surface smoothness, and the structure of uncalcified cartilage. These treatments led to a reduction in Mankin Score by 35% and 53%, respectively. Von Frey assay is commonly used in OA study as a pain outcome by evaluating mechanical allodynia. Using this assay, we observed that OA knees displayed significantly decreased paw withdraw threshold compared to sham knees. However, this OA-related pain was mostly attenuated in Gli1^+^ cell-or purmorphamine-treated knees (Fig. 6D).

**Figure 6.**
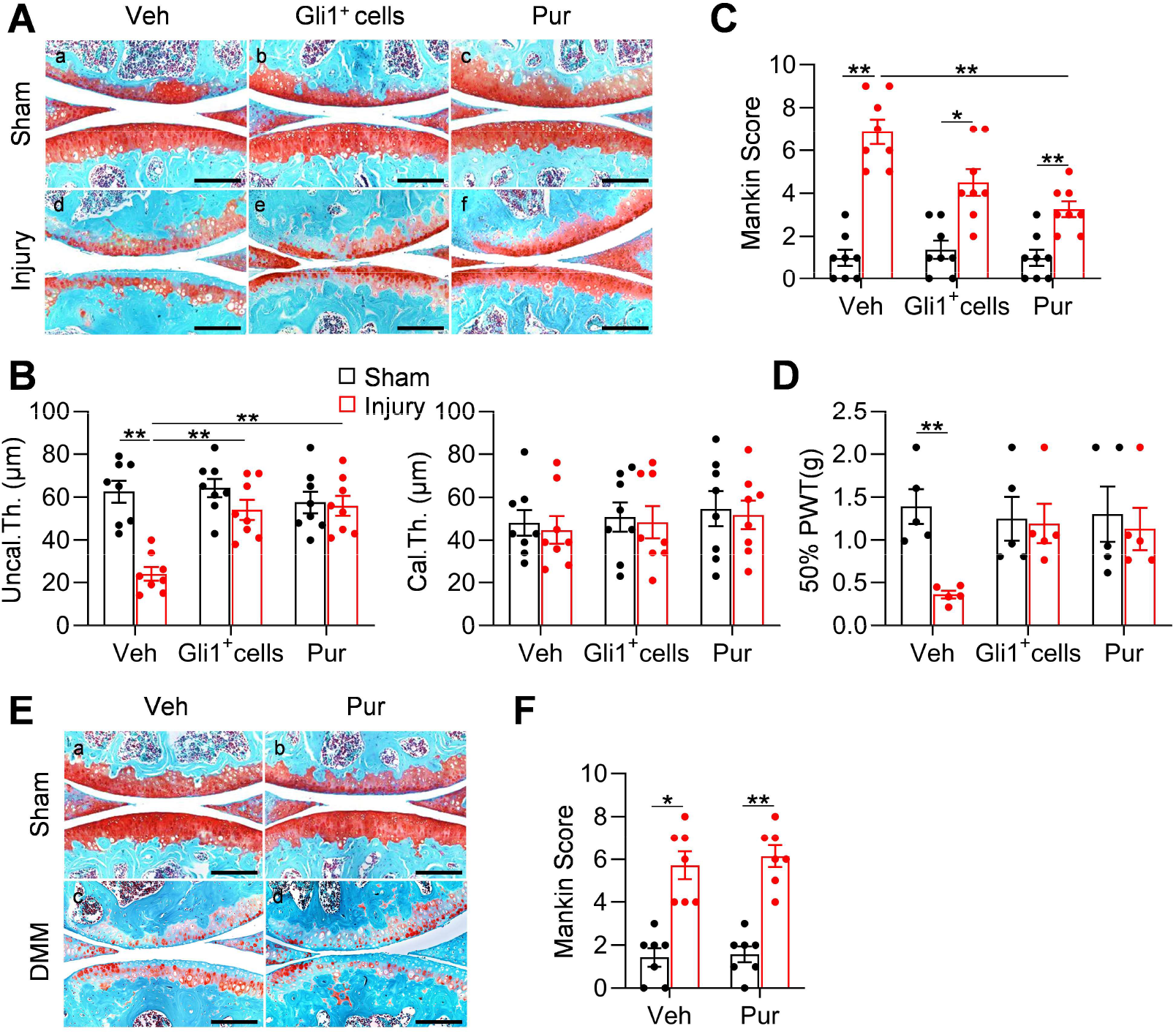
Meniscus repair by enhancing Hh/Gli1 signaling delays OA progression. (**A**) Representative safranin O/fast green staining of sagittal sections of vehicle-, Gli1^+^ cell- and purmorphamine (pur)-treated mouse knee joints at 8 weeks after sham or meniscus injury. Scale bars, 200 μm. (**B**) Average thickness of uncalcified zone (Uncal.Th.), calcified zone (Cal.Th.) of the tibial articular cartilage was quantified. n = 8 mice/group. (**C**) The OA severity was measured by Mankin score. n = 8 mice/group. (**D**) von Frey assay was performed at 8 weeks after injury. PWT: paw withdrawal threshold. n = 5 mice/group. (**E**) Representative safranin O/fast green staining of sagittal sections of vehicle- and purmorphamine-treated mouse knee joints at 8 weeks after sham or DMM surgery. Scale bars, 200 μm. (**F**) The OA severity was measured by Mankin score. n = 7 mice/group. Statistical analysis was performed using two-way ANOVA with Turkey’s post hoc test. Data presented as mean ± s.e.m. *p < 0.05, **p < 0.01, ***p < 0.001.

Hh signaling has also been indicated to play a role in the development of articular cartilage [30] and in OA progression [31]. To exclude the possibility that activating Hh signaling directly affects OA progress, we performed DMM surgery in 3-month-old male *WT* mice and injected purmorphamine into their knee joints right after surgery. Two months later, we observed a similar level of cartilage degeneration in control and treated mice (Fig. 6E, F), suggesting that the effect of Hh signaling on OA development is mediated through meniscus repair but not through directly acting on cartilage. It is worthwhile noting that different from our transient activation approaches, the previous conclusion about the catabolic action of Hh signaling on cartilage is derived from constant modulation of this signaling by genetic approaches.

## Discussion

Previous studies have identified the existence of mesenchymal progenitors in meniscus based on their clonogenic and multi-differentiation activities in culture. However, their in vivo properties and regulatory signals are largely unknown. In this work, by using a lineage tracing line, cell culture, and a meniscus injury model, we demonstrated that Gli1 is not only a mesenchymal progenitor marker in mouse meniscus but that Gli1-labeled cells directly contribute to meniscus development and injury response. Moreover, aging reduces this Gli1^+^ progenitor population in healthy meniscus as well as their expansion after injury, which is consistent with attenuated healing in old mice. On the therapeutic side, the activation of Hh/Gli1 signaling in adult meniscus leads to accelerated meniscus healing process and the delay of OA changes, indicating a protective role of Hh signaling on meniscus against degeneration.

Hh signaling plays a key role during embryonic development and tissue patterning. In long bones, embryonic Gli1^+^ cells give rise to multiple cell types associated with the skeleton and are a major source of osteoblasts in both fetal and postnatal life of the mouse [10]. In another study, embryonic Gli1 lineage cells eventually become the entire mature enthesis by which tendons attach to bone [32]. We also discovered that Gli1^+^ mesenchymal progenitors from neonatal periarticular surfaces are capable of generating mesenchymal lineage cells, including osteoblasts, osteocytes, and adipocytes in the secondary ossification center of long bones [11]. Surprisingly, Gli1^+^ cells do not contribute to the early development of meniscus. They start to appear from 2 weeks of age first in the anterior horn and later in the posterior horn. The different Gli1-labeling patterns in the two horns may reflect the distinct cellular composition of anterior and posterior horns reported previously [33]. Since menisci undergo rapid growth postnatally and Gli1^+^ cells are absent from the meniscus body, our data indicated that there must be other distinct progenitor population(s) contributing to meniscus development. Indeed, we found that Gli1^-^ meniscus cells are also able to proliferate and differentiation in vitro albeit with less activities compared to Gli1^+^ cells.

At the adult stage, we found that Gli1^+^ cells are mainly located at the superficial layer of meniscal horns. They rarely contribute to the inner cells of meniscus probably due to the low turnover of meniscus tissue. However, similar to their counterparts in the periosteum [10] and tendon enthesis [32], they play a major role in tissue regeneration. In our study, we established a meniscus injury model by transection of the anterior horn. A previously reported mouse meniscus injury model (meniscectomy of the anterior horn) revealed almost complete regeneration of meniscus and only subtle cartilage degeneration at 6 weeks post surgery [34]. Compared to that, meniscus repair in our injury model is slow and inefficient with disconnected collagen fibers remaining at the injured site at 3 months post surgery. This prolonged injury causes damage on articular cartilage, leading to moderate OA. Hence, our model is suitable to study the beneficial effects of Hh signaling on meniscus repair and meniscus damage-related OA progression.

Notably, we also observed a quick expansion of synovium enriched with Gli1^+^ cells at the early stage of repair. Therefore, we cannot exclude the possibility that synovial Gli1^+^ cells also contribute to meniscus regeneration. That said, our data showed that Gli1^+^ primary meniscus cells injected into knee joints incorporate into meniscus tissue and accelerate repair, indicating that the endogenous Gli1^+^ meniscus cells are likely responsible for the repair. In addition, we have not investigated the source of Hh protein in meniscus. Since Ihh from the prehypertrophic chondrocytes is known for regulating long bone development through endochondral ossification [35], it is possible that fibrochondrocytes in the deep layer of meniscus produce the Hh signals.

Our lineage tracing and injury studies are based on mouse models. However, rodents are different from human by having bony ossicles in the meniscus horns [36]. To show the clinical relevance of our research, we first demonstrated that porcine meniscus has similar anatomic distribution of Gli1^+^ cells, suggesting a conservation of this patterning between species. While we did not detect Gli1^+^ cells in healthy human meniscus, likely due to the sample being collected from the body rather than the horns, we found similar expansion of Gli1^+^ cells in cell clusters of diseased meniscus, suggesting the translatability of our findings. Our studies, therefore, have uncovered a critical role of Hh/Gli1 signaling in knee meniscus development and regeneration and provide evidence for targeting this pathway as a novel meniscus injury therapy and potentially for preventing OA development.

## Acknowledgments

This study was supported by NIH grants NIH/NIAMS R01AR066098, R21AR074570 (to L.Q.), P30AR069619 (to Penn Center for Musculoskeletal Disorders), and NSF CMMI-1751898 (to LH).

## Author contributions

L.Q., H.S. and Y.W. designed the study. Y.W., H.S., T.G. performed animal experiments. L.Y., L.Z. and W.Y. did all the FACS sorting and flow cytometry analysis. Y.W., H.S., T.G., L.Y., L.Z. and W.Y. performed histology and imaging analysis. Y.W., H.S. and T.G. performed cell culture and qRT-PCR experiments. S.H. performed the meniscal differentiation experiment. L.H., E.K., F.L., M.Z., R.M., S.X.L., Y.Z., and J.A. provided administrative, technical support and consultation. L.Q. and Y.W. wrote the manuscript. L.H., E.K., F.L., R.M., Y.W., S.H., T.G., L.Y., L.Z., S.X.L., M.Z., Y.Z., W.Y., S.H. and J.A. reviewed and revised the manuscript. L.Q. approved the final version.

## Data and materials availability

All data associated with this study are present in the paper and available from the corresponding authors upon reasonable request.

## Supplementary figures

**Fig. S1.**
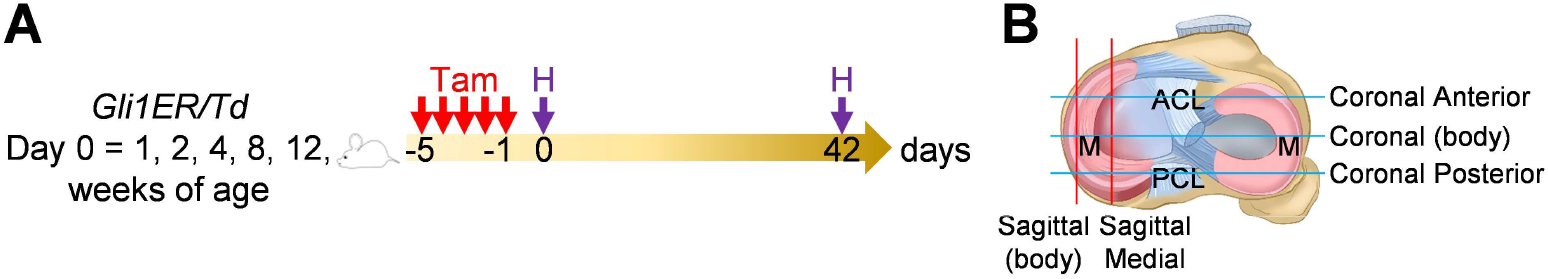
Schematic graph of the study protocol. (**A**) Male *Gli1ER/Td* mice were treated with Tam at 1, 2, 4, 8 and 12 weeks of age and analyzed at 24 h (pulse) or 6 weeks (tracing) after the last Tam dosing. (**B**) Schematic cartoon of meniscus shows sectioning sites. M: meniscus; ACL: anterior cruciate ligament; PCL: posterior cruciate ligament.

**Fig. S2.**
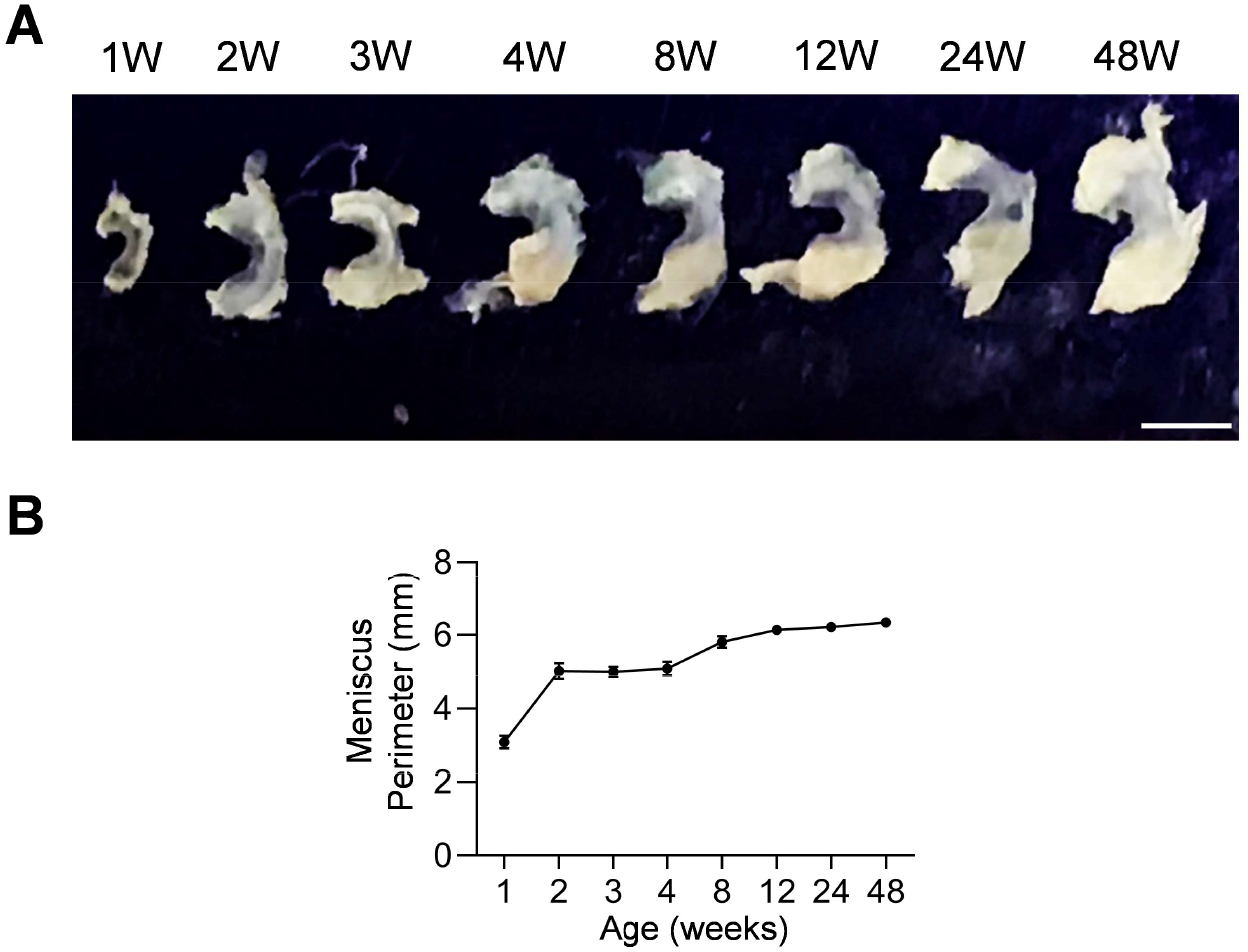
Mouse meniscal morphogenesis during development. (**A**) The morphological overview of meniscus at 1, 2, 3, 4, 8, 12, 24, 48 weeks of age. Scale bars, 1mm. (**B**) The meniscal perimeter was quantified. n = 3 mice/age.

**Fig. S3.**
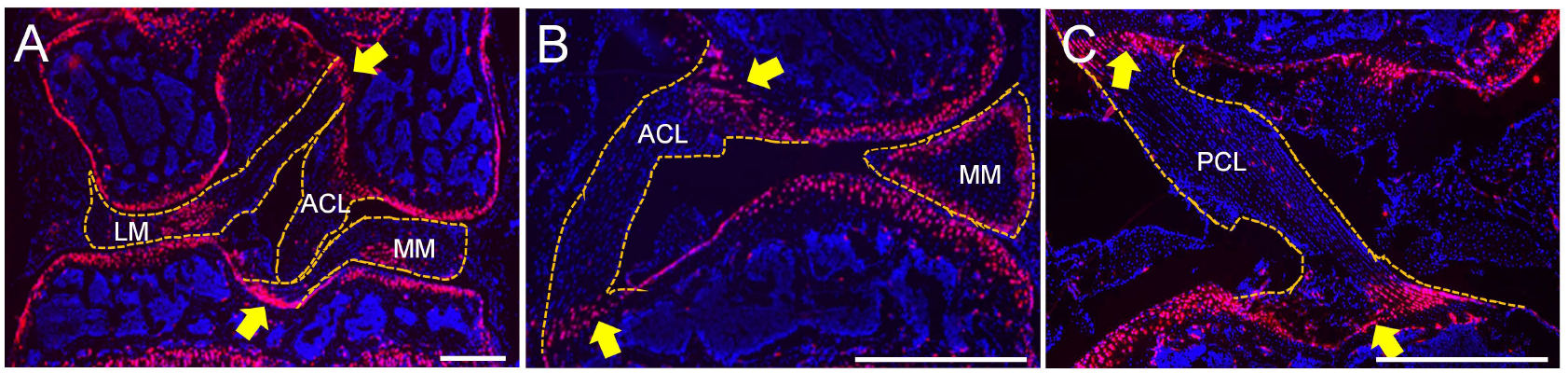
Meniscal enthesis and ligamental enthesis regions in joint are enriched with Gli1 ^+^ cells. *Gli1ER/Td* mice at 12 weeks of age received Tam injections followed by tissue harvest 24 h later. Knees were sectioned to show meniscal enthesis regions (**A**) and ligamental enthesis regions (**B**) and (**C**) within the knee joint. MM: medial meniscus; LM: lateral meniscus; ACL: anterior cruciate ligament; PCL: posterior cruciate ligament. Yellow arrows indicate the enthesis regions.

**Fig. S4.**
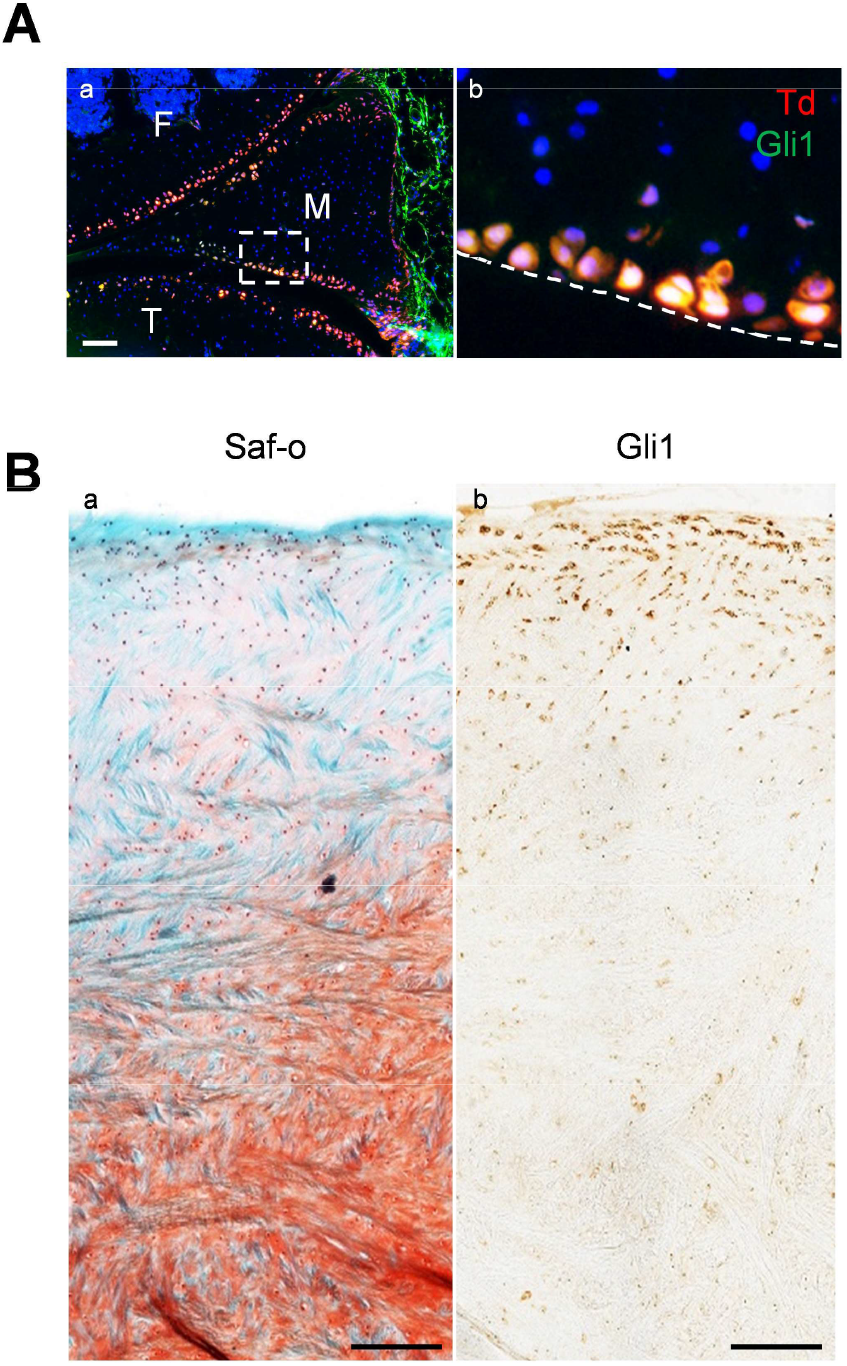
Gli1 labels the superficial zone cells of mouse and mini-pig meniscal horns. (**A**) Immunofluorescence staining of Gli1 (green) on sagittal sections of 12-week-old *Gli1ER/Td* mouse knee joints. Boxed area in a is enlarged in b. Dashed line indicates the surface of meniscus. Scale bars, 200 μm. F: femur; T: tibia; M: meniscus. (**B**) Representative safranin O/fast green staining (left) and immunohistochemistry staining of Gli1 (right) in the horn area of mini-pig meniscus. Scale bars, 200 μm.

**Fig. S5.**
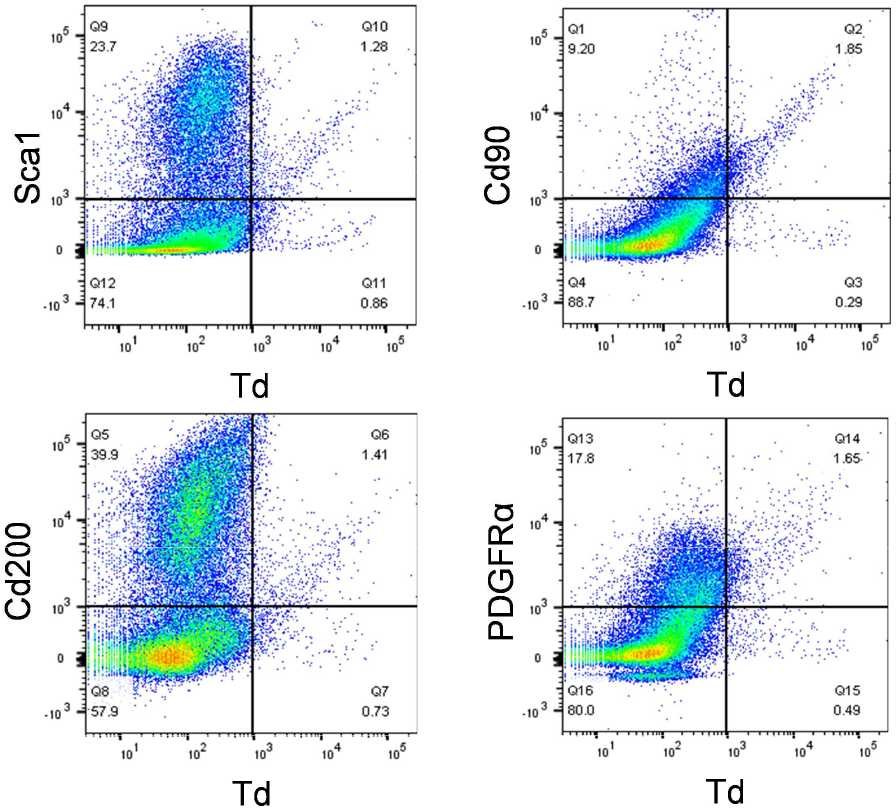
Mesenchymal progenitor markers are enriched in Gli1 ^+^ meniscus cells. Digested meniscus cells from 3-month-old Gli1ER/Td mice were subjected to flow cytometry analysis.

**Fig. S6.**
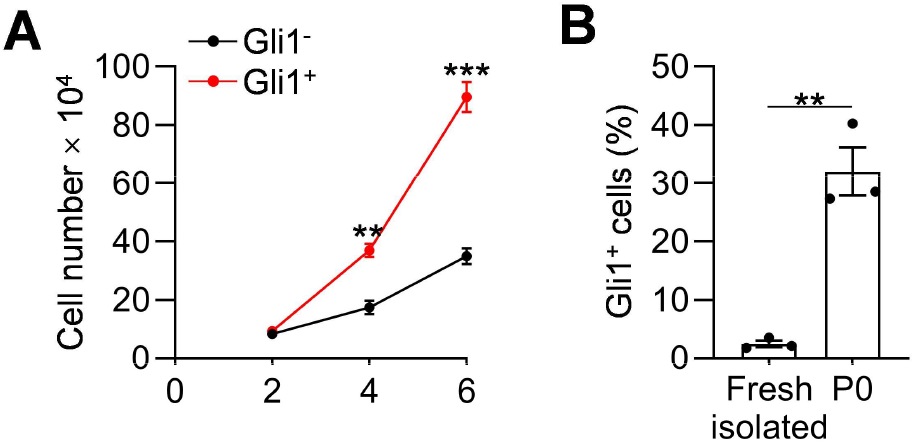
Gli1 ^+^ cells proliferate faster than Gli1 ^-^ cells. (**A**) Sorted Gli1^+^ meniscus cells proliferate faster than Gli1^-^ cells. Cells were seeded at 10,000/well on day 0 and counted every other day. n = 3 independent experiments. (**B**) The percentage of Td (Gli1)^+^ cells from freshly isolated cells and after a 7-day culture was quantified by flow cytometry. n = 3 independent experiments. Statistical analysis was performed using unpaired two-tailed t-test. Data presented as mean ± s.e.m. **p < 0.01, ***p < 0.001.

**Fig. S7.**
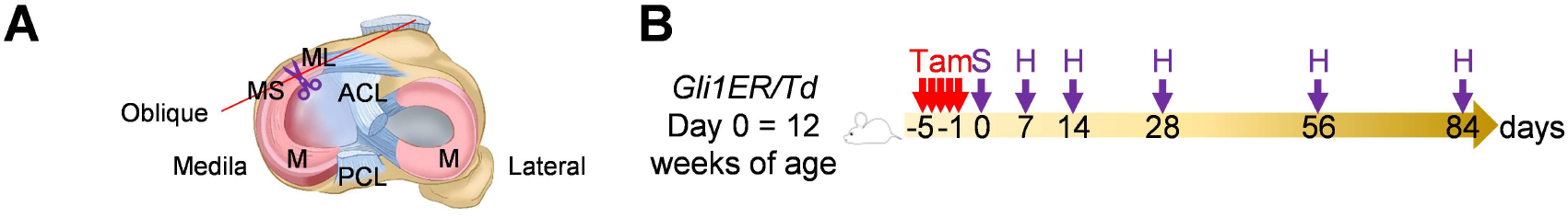
Schematic graph of the study protocol. (**A**) Schematic cartoon of meniscus shows the sectioning site. M: meniscus; MS: meniscus synovial end; ML: meniscus ligamental end. A pair of scissors indicates the transection site. ACL: anterior cruciate ligament; PCL: posterior cruciate ligament. (**B**) Male *Gli1ER/Td* mice received Tam injections (day -5 ∼ -1) and meniscus injury (day 0) at 12 weeks of age. Knee joints were harvested at 1, 2, 4, 8, 12 weeks after injury.

**Fig. S8.**
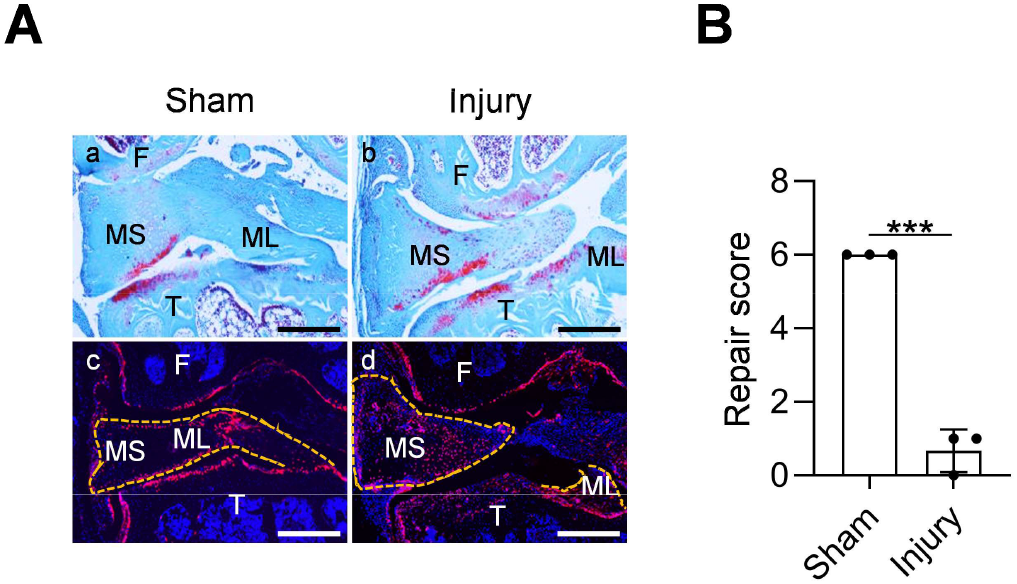
Aging diminishes Gli1 ^+^ cell expansion and the repair ability of meniscus. (**A**) Representative safranin O/fast green staining (top) and fluorescence images (bottom) of aged mouse knee joints at 4 weeks after sham or meniscus injury. *Gli1ER/Td* mice at 12 months of age received Tam followed by meniscus injury. Dashed lines outline the meniscus. Scale bars, 200 μm. F: femur; T: tibia; MS: meniscus synovial end; ML: meniscus ligamental end; Red: Td; Blue: DAPI. (**B**) Repair score was quantified. n = 3 mice/group. Statistical analysis was performed using unpaired two-tailed t-test. Data presented as mean ± s.e.m. ***p < 0.001.

**Fig. S9.**
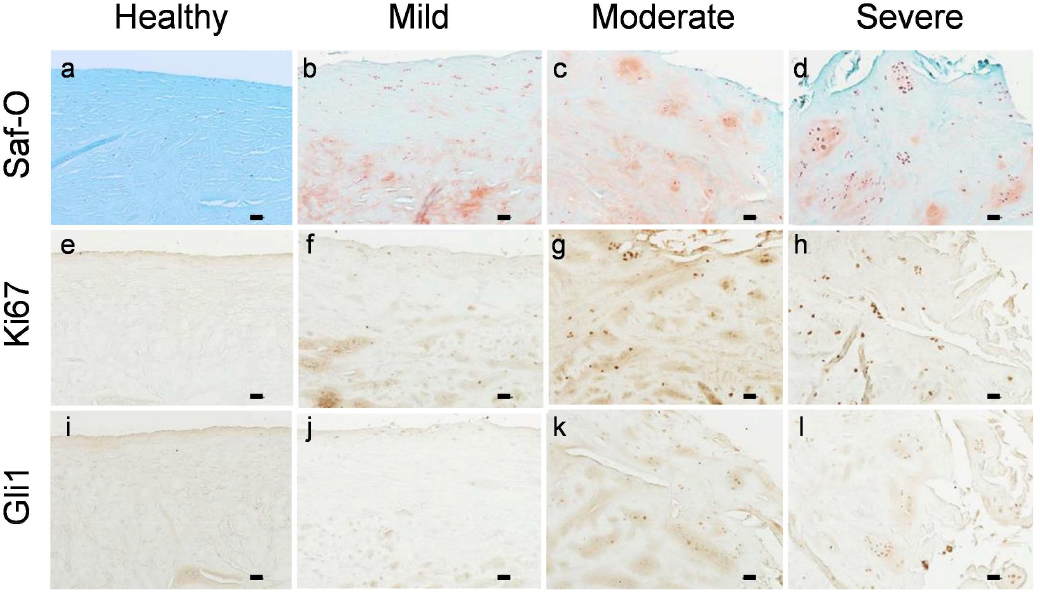
Gli1 ^+^ cells appear in proliferative cell clusters of degenerated human meniscus. Representative safranin O/fast green staining (top) and immunohistochemistry staining of Ki67 (middle) and Gli1 (bottom) in human meniscus tissues at different degenerative stages. n = 3 samples/stage. Scale bars, 200 μm.

**Table S1.**
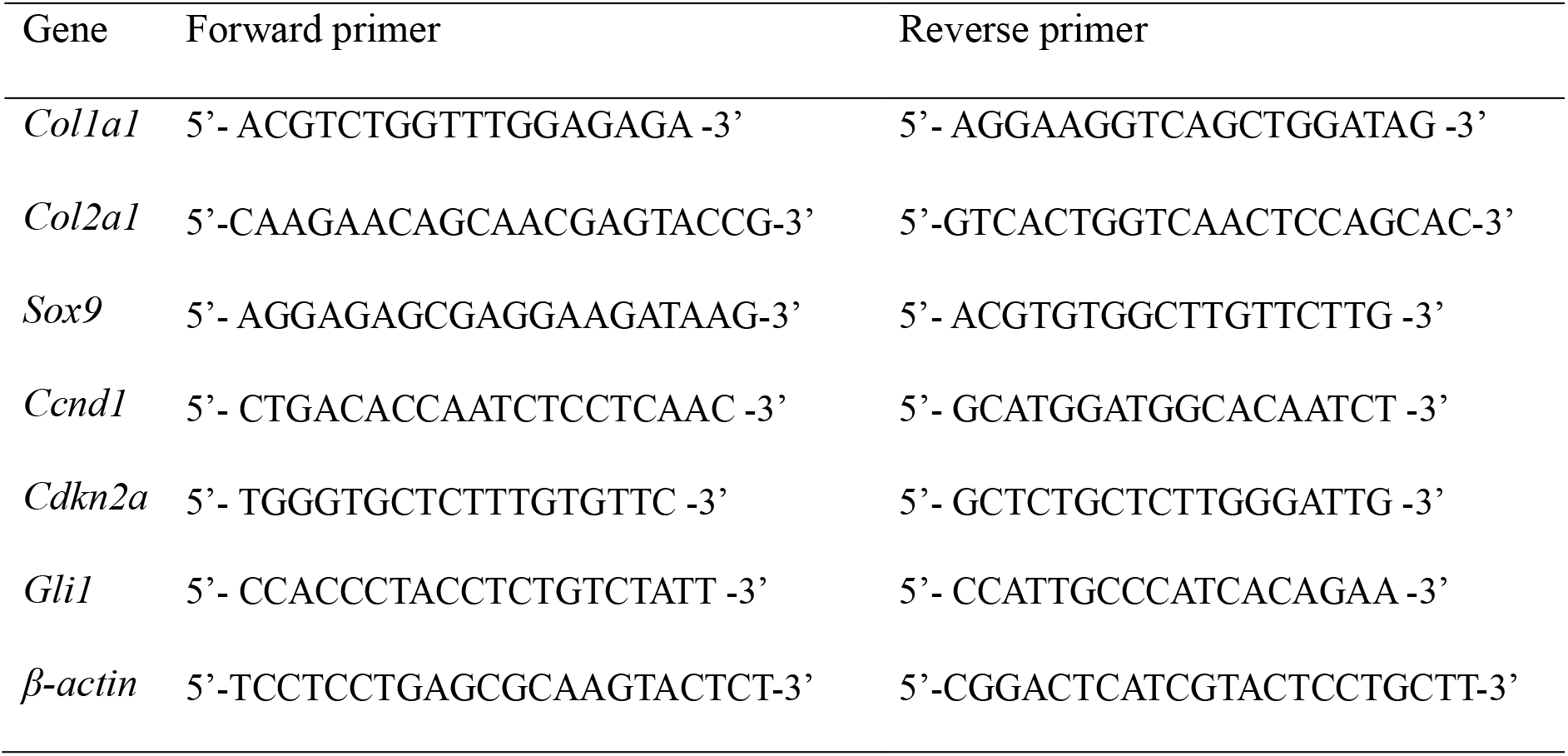
Mouse real-time PCR primer sequences.

